# Review: Population Structure in Genetic Studies: Confounding Factors and Mixed Models

**DOI:** 10.1101/092106

**Authors:** Lana S. Martin, Eleazar Eskin

## Abstract

A genome-wide association study (GWAS) seeks to identify genetic variants that contribute to the development and progression of a specific disease. Over the past 10 years, new approaches using mixed models have emerged to mitigate the deleterious effects of population structure and relatedness in association studies. However, developing GWAS techniques to effectively test for association while correcting for population structure is a computational and statistical challenge. Using laboratory mouse strains as an example, our review characterizes the problem of population structure in association studies and describes how it can cause false positive associations. We then motivate mixed models in the context of unmodeled factors.

## Introduction

Genetics studies have identified thousands of variants implicated in dozens of common human diseases (Manolio et al. 2009; Purcell et al. 2009; Stram 2013; Yang et al. 2010). These variants are locations in the human genome where genetic content differ among individuals in a population. A genome-wide association study (GWAS) seeks to identify genetic variants that contribute to the development and progression of a specific disease.

Association studies discover these genetic factors by correlating an individual’s genetic variation with a disease status or disease-related trait. At the genome-wide scale, association studies typically focus on statistical relationships between single-nucleotide polymorphisms (SNPs) and disease traits. SNPs are the most common genetic variants underlying susceptibility to disease, and associated SNPs are considered to mark the region of a human genome that influences disease risk. A GWAS identifies a SNP as a significant, and therefore *associated,* variant when the specific genome sequence at the SNP is correlated with a disease trait or disease status. For example, a GWAS study may find that individuals with a specific sequence (or allele) at a SNP have higher blood pressure on average than individuals with a different sequence at the SNP. If a SNP has a significant correlation with a trait or disease status, the association study suggests that presence of the particular variant may increase an individual’s risk for disease.

Typical analytical strategies for performing association studies rely on standard regression techniques, which assume the data have an identically and independently distributed (*i.i.d.*) property. If data have *iid*, all variables are mutually independent since each random variable shares the same probability distribution with other variables. Association study methodology was originally designed for populations comprised of unrelated individuals, and standard approaches assume this property is true (Risch and Merikangas 1996). However, the big genomic datasets available today inevitably contain distantly related individuals. This genetic relatedness prevents standard association studies from correctly identifying the causal variants and induces identification of many false positive associations (or *spurious associations*).

Two types of relatedness may produce high rates of false positive associations: population structure and cryptic relatedness. *Population structure* refers to different ancestry among individuals in a study. *Cryptic relatedness* exists when some individuals are closely related, but this shared ancestry is unknown to the investigators. Large (n=>5000) population cohorts inevitably contain individuals who have common ancestry from different populations. In either case, individuals who share ancestry are more related than individuals from different ancestries. These ancestry differences induce a self-organizing population structure effect, which causes the statistical methodology to assign strong association signals to variants that are not actually causal for the trait or disease. In many cases, applying standard association study techniques to population cohorts with population structure produces a high rate of false positive associations. These associations may appear to be significant, but they are driven by the cohort’s relatedness rather than variants that truly affect trait or disease risk.

Developing GWAS techniques to effectively test for association while correcting for population structure is a computational and statistical challenge. This challenge is relevant to human association studies as well as genetic studies in any organism, including model organisms such as mice. Mouse studies are widely used to study human disease and, because the particular history of laboratory mouse strains induces complex patterns of genetic relatedness that can cause false positives in association studies.

Over the past 10 years, new approaches using mixed models have emerged to mitigate the deleterious effects of population structure and relatedness in association studies (Zhou and Stephens 2012; Kang et al. 2008, 2010; Listgarten et al. 2012). These approaches were originally developed in the context of mouse studies and later applied to human studies. In this review, we explicitly characterize population structure as a confounding factor in order to explore the root cause of false positives in association studies. We trace the development of these methods in mouse studies and describe how these methods were adapted to human studies, particularly where they are applied to correct for population structure in large-scale genomic datasets.

### Standard Genome Wide Association Studies (GWAS)

Genetic association studies attempt to identify single-nucleotide polymorphisms (SNPs) that are responsible for differences in trait or phenotype values within an individual. A SNP is a single position in the human genome sequence where individuals in the population have different genetic content. These differing forms of the same gene are referred to as alleles. SNPs are the most common form of genetic variation, and almost all common SNPs have two alleles.

SNPs are ideal targets for association testing, because they are the most common form of genetic variants. Their high level of prevalence means that they are often correlated with other forms of variation. To conduct a typical single-SNP test, we first collect genetic information at the SNP in a set of individuals (referred to as genotypes). Next, we measure the association (or correlation) with the trait values (or phenotypes) of the individuals (see Figure 1a). In this Figure, it is intuitively clear that the first SNP appears to be associated, but the second SNP does not appear to be associated.

In order to evaluate if the association between a SNP and a phenotype is statistically significant, we can use the collected data to test two hypotheses. The null hypothesis assumes a model where the SNP does not affect the phenotype (see Figure 1b). In this hypothesis, the phenotypes (*y*) are only affected by the population mean (*µ*) and the environment (*e*). Unless data indicate otherwise, we assume that the null hypothesis is true and the SNP does not influence the phenotype (i.e., does not affect the individual’s disease risk).

An alternative hypothesis provides a model of the SNP being significantly associated with the phenotype (see Figure 1c). In this case, the phenotypes (*y*) are affected not only by the population mean (*µ*) and environment (*e*), but they are also affected by the genotype (*x*). In other words, presence of the SNP suggests an individual is likely to have the trait or disease risk. Here, the quantitative measurement of strength that the genotype has on the phenotype is referred to as the effect size (*β*). If the effect size (*β*) is equal to 0, we consider the two models equivalent. The SNP is determined to be significantly associated with the phenotype when the data fits the alternative hypothesis beyond a specific threshold.

**Figure 1.**
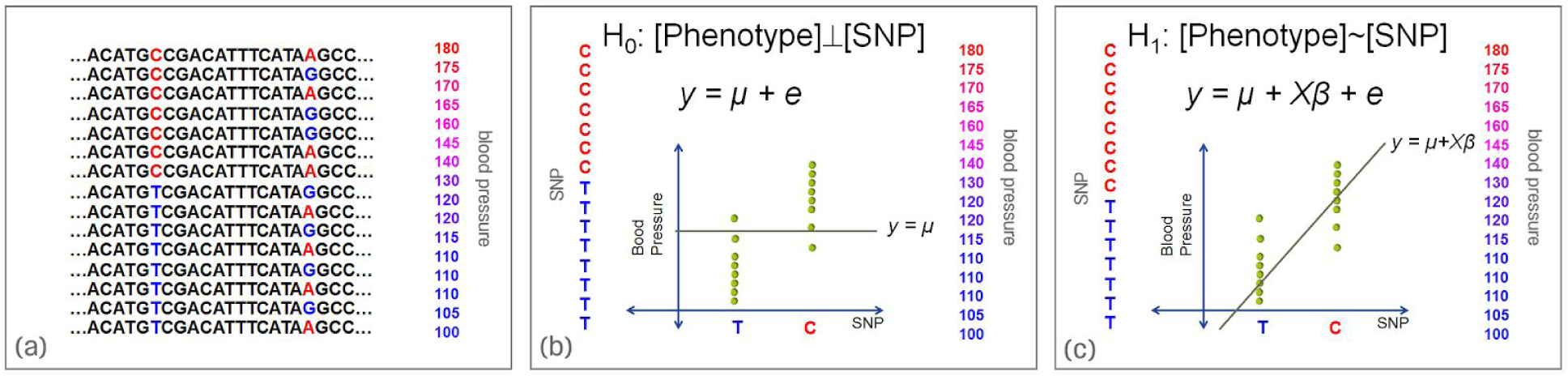
Standard genetic association study applied to human blood pressure data. (a) The left SNP appears to be more strongly associated with blood pressure than the right SNP. (b) We test two hypotheses against each other to evaluate whether the association between a SNP and a phenotype is statistically significant. By default, a null hypothesis assumes that the SNP does not affect the phenotype. (c) If the data fits the alternative hypothesis beyond a certain threshold, the SNP is described as significantly associated with the phenotype.

We mathematically express the null and the alternative hypotheses in order to perform a single-SNP test. We denote the kth genotype of the jth individual *g*_*j*__*k*_ where the genotype is in the set {0,1,2}, which is the number of copies of the kth variant that the jth individual has on their two chromosomes. Here, a “0” denotes the genotype that does not contain the variant in either chromosome, while a “1” or “2” denotes the genotype presence in one or two of the chromosomes, respectively. In order to simplify the equations for association studies, we standardize the genotypes by subtracting the population mean and dividing by the variance. The frequency of a variant in the population is denoted as *p*_*k*_, which is the average genotype frequency in the population. The standardized genotypes can be expressed as

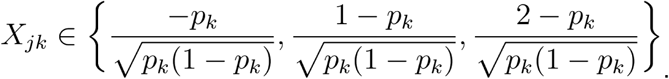

Once we have calculated the standardized genotypes, a typical single-SNP test can be used to identify variants associated with traits. A standard regression technique estimates the relationship among variables, including a dependent variable (*y*), any independent variables (*x*), and unknown variables (*β*). Using regression, these simple linear models can correlate the genetic variation with the trait, allowing us to assess whether the data best fits the null or alternative hypothesis.

The equation

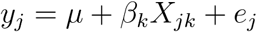

models the phenotype for a single individual *j* in the study. Here, the effect of the variant on the phenotype is *β*_*k*_, the model mean is *µ*, and the contribution of the environment on the phenotype is *e*_*j*_. The environment’s effect on a phenotype for an individual *j* (*e*_*j*_) is assumed to be normally distributed with variance 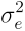
 denoted as 
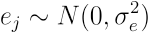

The equation above describes the relationship between the genotype and phenotype of just one individual. We can use vector notation to represent all of the individuals in the dataset and produce the model

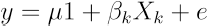

with the phenotypes of all of the individuals in the dataset denoted as column vector *y*, a column containing the genotypes for the ith variant in the population denoted as *x*_*i*_, and a vector containing the environments denoted as e. 1 is a column vector of 1s. We draw the random vector *e* from the distribution
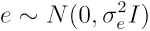
. We note that each element of is independent of the others; hence, the variance-covariance matrix is a diagonal matrix 
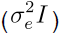
.

We can write the distribution of *y* using

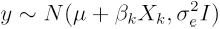

where *I* is the identity matrix.

Using the observed data (such as the example in Figure 1), we can estimate the values of the population mean and the effect of the true variant by using the following equations:

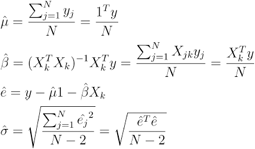

The reason the equations are so simple is because the genotypes are standardized. The resulting value is the association between an SNP and phenotype. We can then test the significance of this association by using the following statistic:

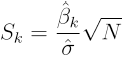

This statistic is normally distributed with a mean that depends on the effect of the SNP on the trait, the environmental variance, and the number of individuals. The variance of the statistic is 1. We can write the distribution of the statistic as

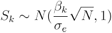

If the SNP does not have an effect on the trait, the statistic will follow the null distribution

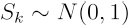

which is a standard normal distribution. We can then use this null distribution to determine whether the association is significant. This statistic is considered significant with a significance level of *α_s_* if

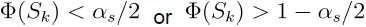

in which case the variant is considered to be associated (see Figure 2). We use the notation *α_s_* to denote the significance level that we need to achieve at any SNP, which in human studies is typically 5*10^−8^.

**Figure 2.**
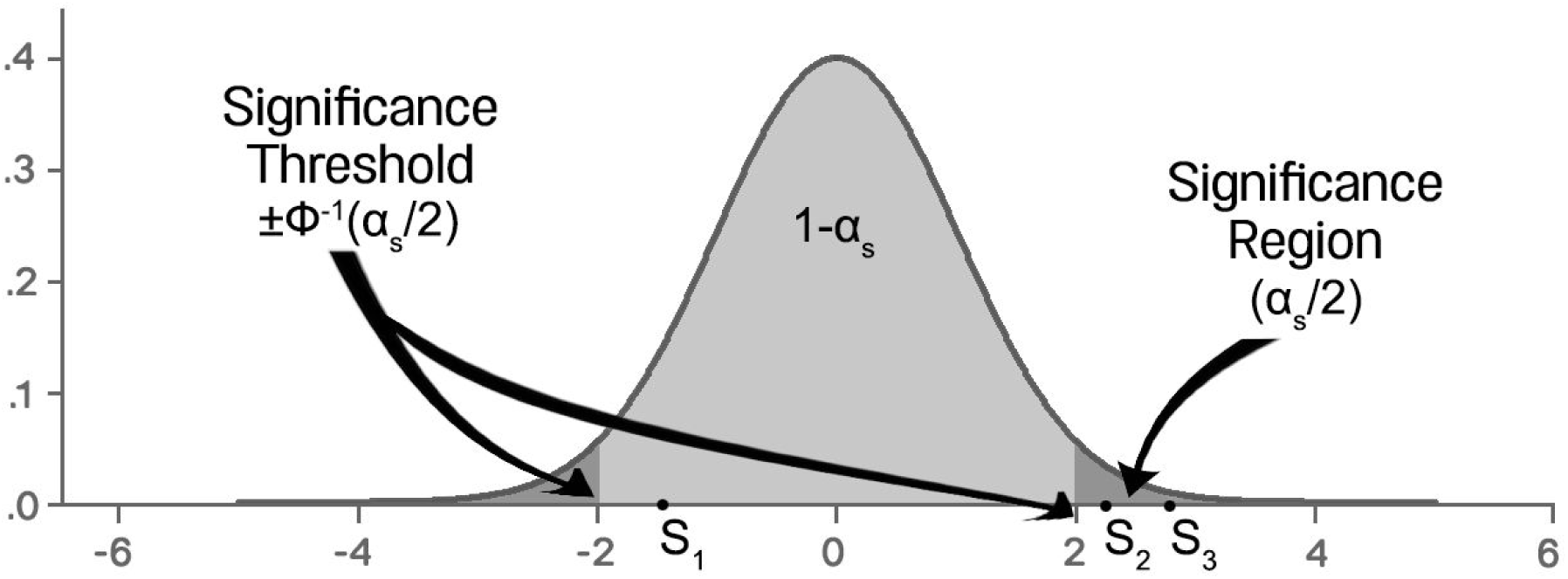
Significance testing in association studies. The null distribution is the standard normal distribution and the expected distribution of the association statistics under the assumption that the effect size is 0. Each variant’s association statistic in equation (3) is computed, and its significance is evaluated using the null distribution. If the statistic falls in the significance region of the distribution, the variant is declared associated. In this example, S1 is *not* significant, while S2 and S3 *are* significant. The exact location of the threshold is defined as the location on the x-axis where the tail probability area equals the significance threshold (S). This is denoted using the quantile of the standard normal □^*ϕ*^−1^(*x*)^.

The p-value of the association is the tail probability area beyond the observed statistic, and the p-value can be computed using
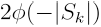
 If the SNP does not affect the trait, the statistic will come from the null distribution. In this case, the p-values will be uniformly distributed between 0 and 1.

## True Genetic Model

Assuming *iid*, the single-SNP test will tell us if a SNP is responsible for the differences we observe in an individual’s trait or phenotype expression values. However, this simple linear model is an unrealistic model for identifying variants associated with traits in today’s large genomic datasets that contain a high degree of relatedness. In real populations, the true effect of a single SNP is influenced by multiple variants that are affecting the trait. A ‘hypothetical’ true genetic model takes into account the effect of all SNPs on the trait.

Here, the vector notation

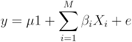

models the phenotypes of all the individuals in the dataset denoted as column vector *y*. Again, the effect of the *i* th variant on the phenotype is *β_i_*, the mean is *μ*, and the contribution of the environment on the phenotype is denoted by *e*. Here, the number of variants is *M*.

The true genetic model takes into account the true effect of all SNPs, including the effect of the SNP being tested for association with a trait. When testing SNP *k*, we are using equation (1) the actual data is generated from

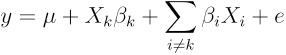

In applying the simple linear model to data, we observe a mismatch between the model used for testing and the assumed underlying generative model. Here, any term that is missing in the testing model when compared to the generative model is called an *unmodeled factor*. The unmodeled factor is exactly 
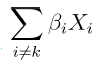
.

In this case, the unmodeled factor is the effect of variants in a genome other than the variant being tested. This factor can significantly affect the results of an association study. If the individuals in the study are related to each other, the unmodeled factor may produce a high rate of false positive associations.

In an association study, relatedness among individuals is referred to as population structure. Over the past few years, there have been many methods which have been developed to mitigate the effect of population structure in association studies. One of the most commonly utilized approaches today, mixed models, originally became popularized in mouse studies and is now the standard approach for analyzing human GWAS studies. In this review, we motivate the problem of population structure in association studies utilizing laboratory mouse strains and explain how it can cause false positives associations. We then motivate mixed models in the context of unmodeled factors.

### An example of Population Structure Confounding from Mouse Genetics

The importance of controlling for population structure is evident in genetic mapping of inbred mouse strains. Mice strains pose particular problems that mixed models are developed to solve, and the basic ideas behind mixed models can be clearly demonstrated with mice genetics. Today’s classical inbred laboratory mouse strains descend from a relatively small number of genetic founders (mostly fancy mice originally kept as pets) and are characterized by several population bottlenecks (Frazer et al. 2007; Yang et al. 2007).

A second group of laboratory strains are referred to as “wild-derived” strains. These strains are mouse strains captured from wild and inbred mice that were never kept as pets. Wild-derived strains do not share the population history of classical laboratory strains. A simple way to visualize the relationship between multiple ancestral groups and traits in the mouse genome is with a phylogenetic tree that can be computed from the genetic information (Figure 3). This tree visualizes the genetic relationships between 32 classical inbred strains and 6 wild derived strains, using genetic variant information at 140,000 SNPs for each strain.

We observe that the two groups are close to each other in the phylogeny and are separated by a long branch length (denoted with a dotted line). This branch represents the many genetic differences between the groups. We also have measurements for body weight and liver weight for each of the two strains. Not surprisingly, the body weights of the classical strains are much larger than the body weights of the wild derived strains (Figure 4). Different selective pressures on the two groups, including environmental fitness (wild-derived) and human selection (laboratory), produced these differences in population genetics.

**Figure 3.**
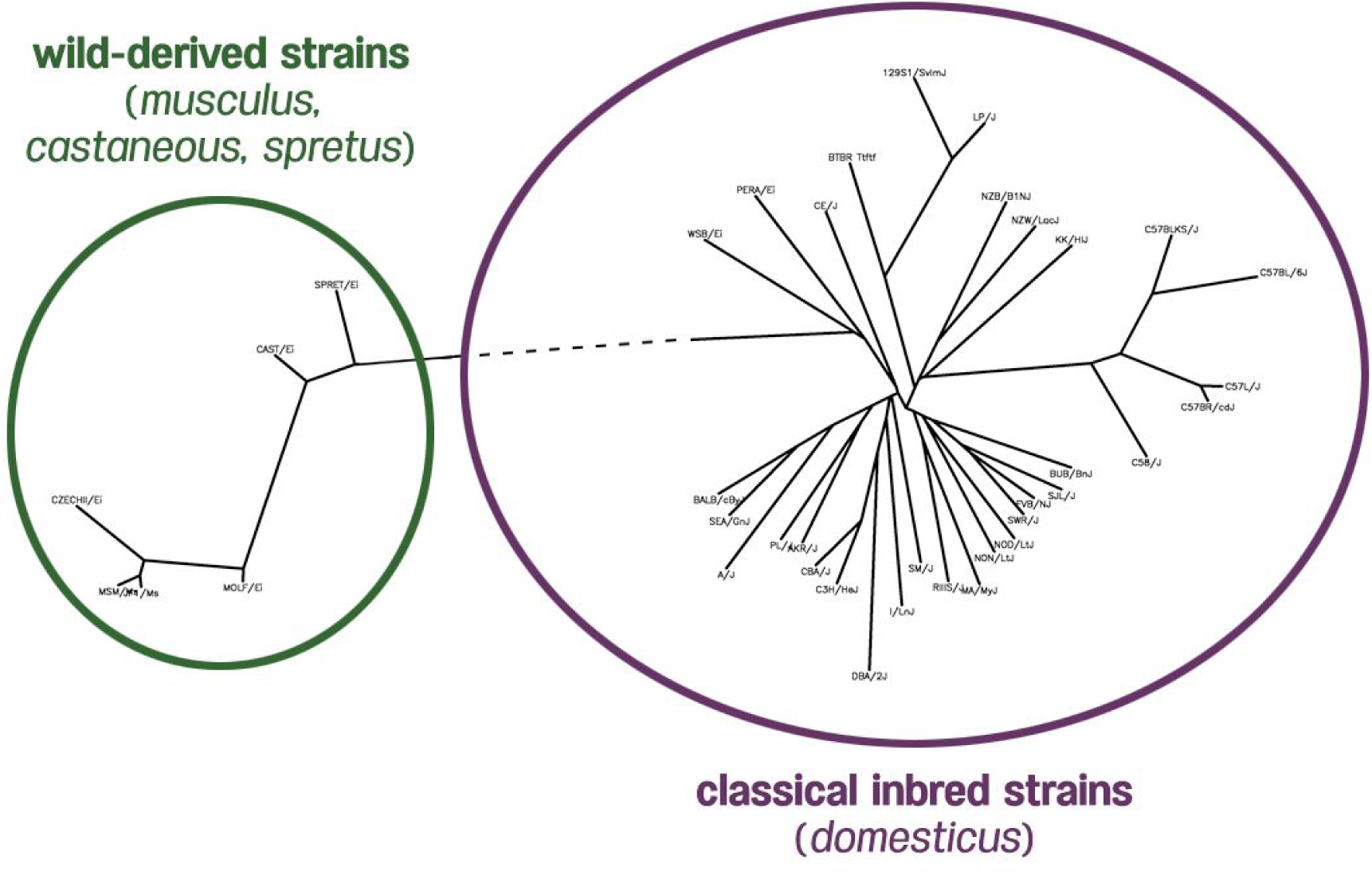
Phylogeny of 38 inbred mouse strains using 140,000 mouse HapMap SNPs. Green strains represent wild-derived non-domesticus mice, and purple strains represent classical inbred mice.

**Figure 4.**
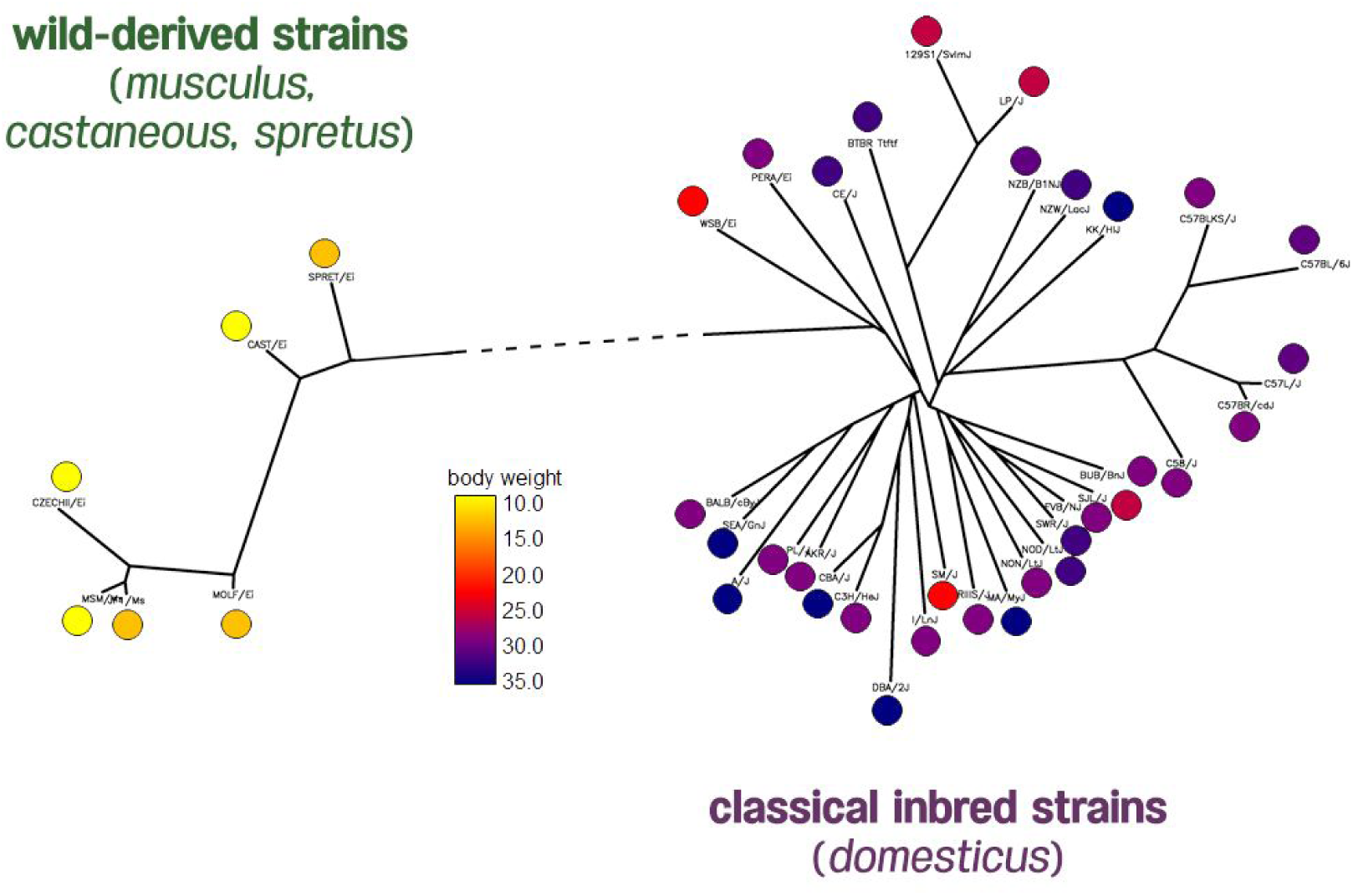
Body weight phenotypes of 38 inbred mouse strains from the Mouse Phenome Database generated by The Jackson Laboratory. The distribution of mice body weights shows the two clades of mice having very different body weights.

In order to identify which genetic variants are associated with body weight, we applied the linear model described above to the 140,000 SNPs from this dataset. In general, we expect association study results to indicate very few significant associations between particular SNPs and a trait. One common way to visualize the results of an association study is with a Manhattan plot. In a Manhattan plot, the mouse genome is plotted against the x-axis, and the measure of significance of correlation between the genome and trait is plotted against the y-axis. Each red spike represents a SNP at a particular genomic position, and the height of the spike represents the strength of the association. The green horizontal line represents the significance threshold. Any SNP that crosses this line is considered a significant association.

We expect to observe a Manhattan plot similar to the one in Figure 5, where a number of SNPs affect the phenotypes. Thus, we would observe signals that cross the threshold at a few locations in the genome, but most of the SNPs will not be associated with the phenotype.

Another way to visualize the results of an association study is with a cumulative p-value distribution plot (b) and a quantile-quantile (Q-Q) plot (c). These plots are graphical techniques for determining whether multiple datasets come from populations with common distribution.

Here, the cumulative p-value distribution plot shows the quantiles of the p-values, which assess the probable significance of association between a genotype and trait; the Q-Q plot shows the distribution of the same data log-transformed.

Since we expect most SNPs not be to associated, most of the statistics will be coming from the null distribution. Thus, most of the p-values will be uniformly distributed between 0 and 1. Typically, only a small fraction of the SNPs have signals stronger than expected at the tail of the distribution. This results in a cumulative p-value distribution that is close to the diagonal line (Figure 5b) and a Q-Q plot that follows the line for the beginning of the curve (as shown in Figure 5c). As shown in Figure 5, we would expect that the median p-value would be close to 0.5.

**Figure 5.**
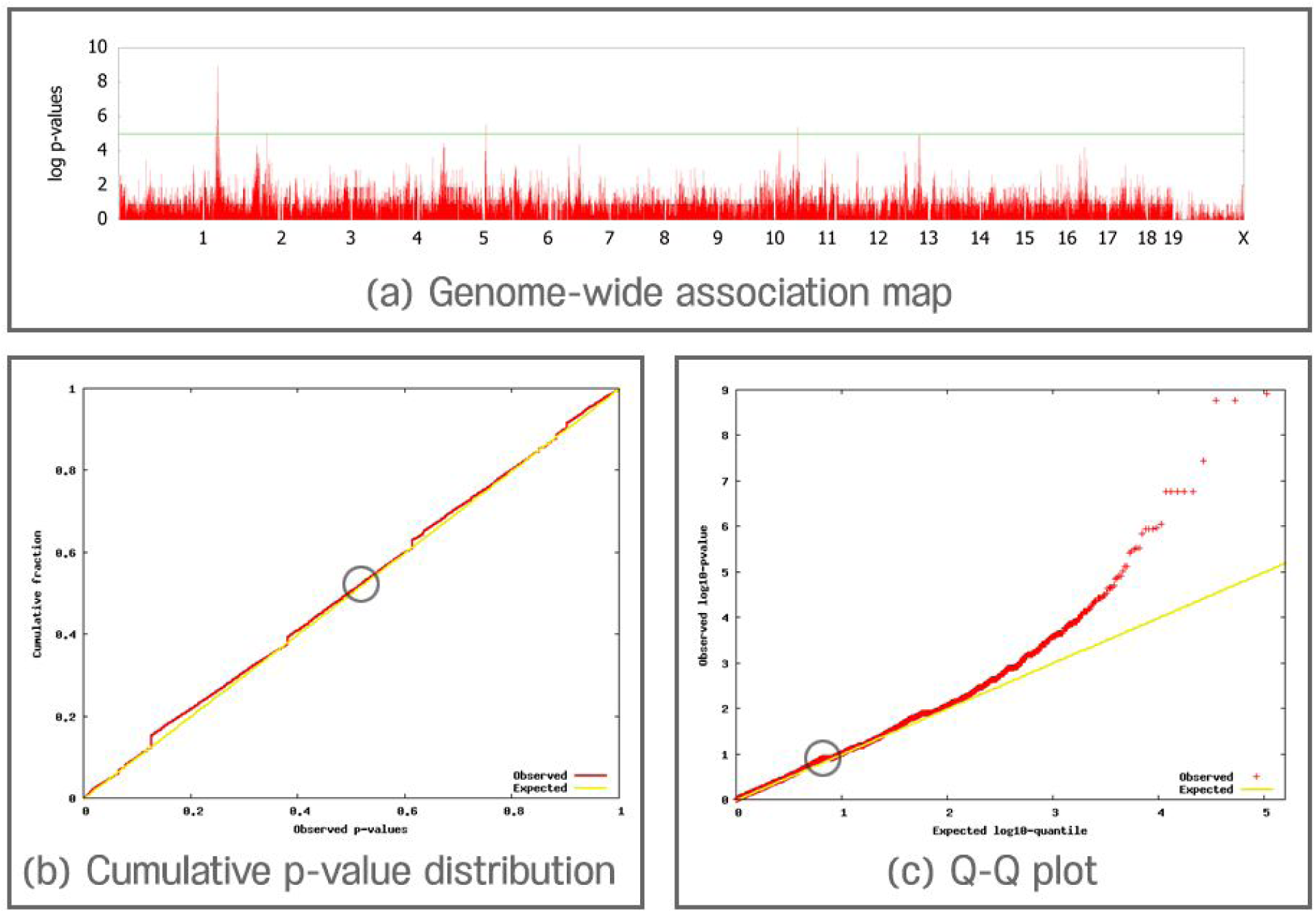
Expected distribution of p-values in a typical (a) Manhattan plot, (b) genome-wide association plot, and (c) Q-Q plot. Circles in (b) and (c) denote where the median p-value (red line) falls on the graph in comparison to the expected median p-value (yellow line). Here, the median falls close to 0.5, suggesting population structure is not affecting association results or has been corrected for in the model.

However, when we applied standard linear models to the inbred mouse dataset, we observed strong signals in many locations in the genome (Figure 6a). The cumulative p-value distribution and the Q-Q plots are shown in Figure 6b and 6c. In our results, we observe that nearly 50% of the SNPs are significantly associated with the phenotype. There are far more significant associations (red line) than expected associations (yellow line).

**Figure 6.**
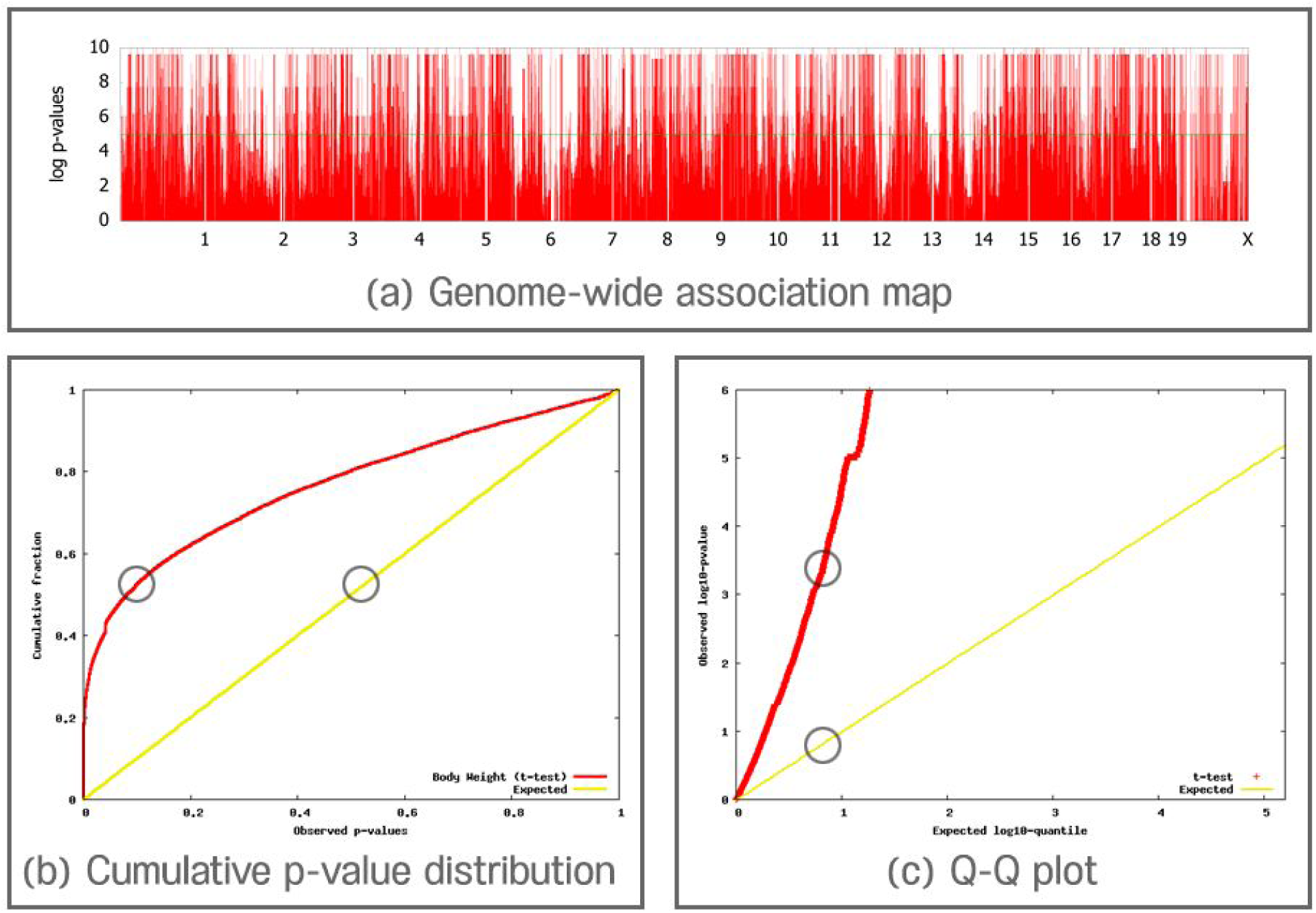
Observed distribution in a (a) Manhattan plot, (b) genome-wide association plot, and (c) Q-Q plot. Circles in (b) and (c) indicate where the median p-value falls on the plot compared to where it is expected. The genomic control factor is computed by considering the observed median p-value compared to the expected. Here, there is a substantial deviation between the red and yellow lines due to inflation of false positive associations for the body weight phenotype.

### Why We Observe False Positives in Mouse Genetic Studies

We can explain why we observe the excess amount of strong association by examining the data for one of the red peaks from the Manhattan plot (Figure 6a) in Figure 7a. Here, the big circles are body weight values, and the small circles are genome-wide SNPs. When we look at the distribution of body weight values and SNPs, it appears that green SNPs correspond to mice with small body weight, while pink SNPs correspond to mice with heavy body weight. Clearly there is a very strong correlation between the SNP and the trait of body weight; it is no surprise that we observe a very significant p-value.

However, if we lay the phylogenetic tree over the pattern of SNPs and body weight values (Figure 7b), we see that the separation of the population into classical and wild derived strains is strongly correlated with the body weight. Here, the SNP differentiates these two groups. The length of each branch in the tree corresponds to the amount of genetic differences between the two groups separated by the branch. The long branch length between the classical and wild strains indicates that many SNPs are dominant in one group and each has a strong signal. This correlation between strains and SNPs causes the large amount of observed associations.

Clearly there are genetic differences between these two groups that affect body weight, but not every genetic difference between the two groups affects body weight. However, the simple linear model will associate every SNP that separates these two groups with body weight. Thus, most of the associations that we observe are for SNPs that are not actually affecting body weight. These associations are referred to as spurious associations.

**Figure 7.**
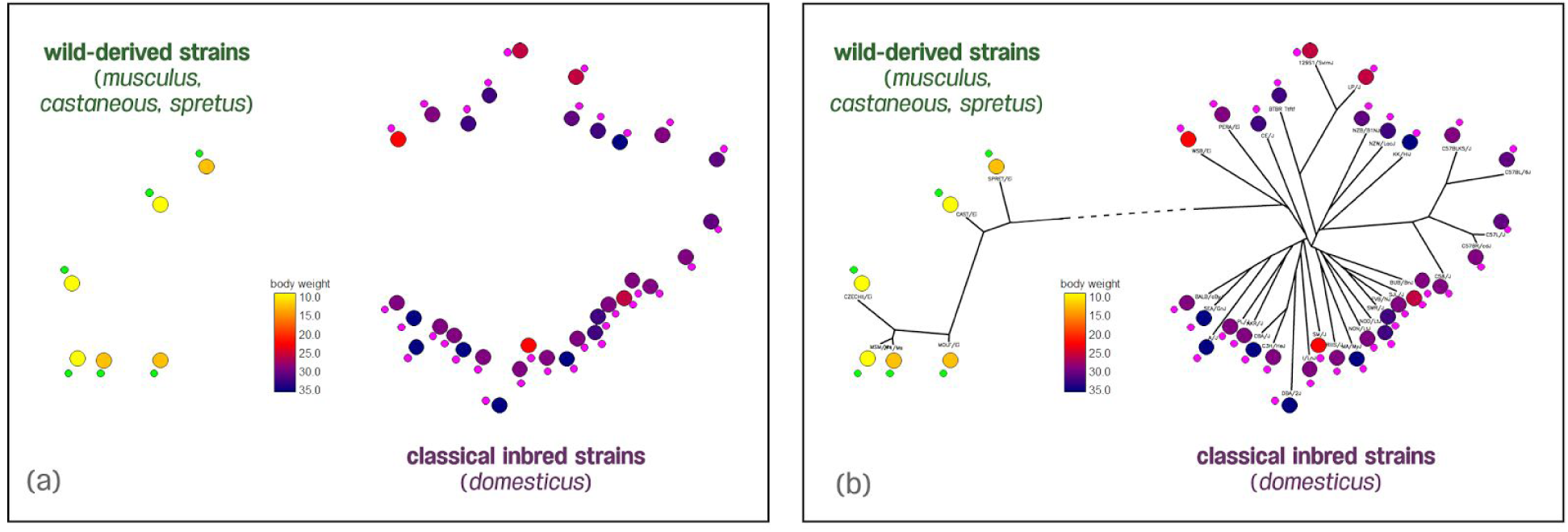
Body weight phenotypes of 38 inbred mouse strains from the Mouse Phenome Database. (a) When we look at only SNP distribution, it appears that green SNPs correspond to small mice, while purple SNPs correspond to large mice. (b) When we look at SNP distribution and phylogeny together, we see that many SNPs segregate the two clades due to a long, shared breeding history.

Another way to understand the effect of population structure on association is through graphical models. We consider SNPs and traits in Figure 8a. Typically, we perform an association test on a SNP. Observation of an association gives evidence that the SNP affects the trait. On the other hand, if we don’t observe an association, this suggests that either the SNP does not affect the trait, or that the effect is too small for our study to detect. However, if genetic differences between groups are present (Figure 8b), shared histories will produce many SNPs directly correlated with population structure (straight dark line). In addition, phenotypes, such as body weight, are also highly correlated with the population structure (straight dark line). This will induce correlation between many SNPs and the phenotype (dotted line) including, but not limited to, the SNPs that are actually responsible for variants.

This phenomenon of association due to relatedness is exactly related to Equation (3). Here, the genetic history shared between mouse strains is the unmodeled factor 
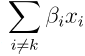

Since the shared genetic history is missing from the testing model, we consider population structure the unmodeled factor.

### Using Mixed Model Methods in Mouse Association Studies

We have shown that population structure can bias association study results. Our mouse examples show that we must correct for population structure in order to accurately identify specific genetic variants involved in disease risk. Several challenges presently limit usefulness of genome association studies for implicating genetic variants. First, unmodeled factors are not known and cannot be accounted for in computational methods that match traits with phenotypes. Second, we do not know the exact ways that unmodeled factors interact with population structure to bias output. Finally, many studies ignore dependency among these unmodeled factors.

**Figure 8.**
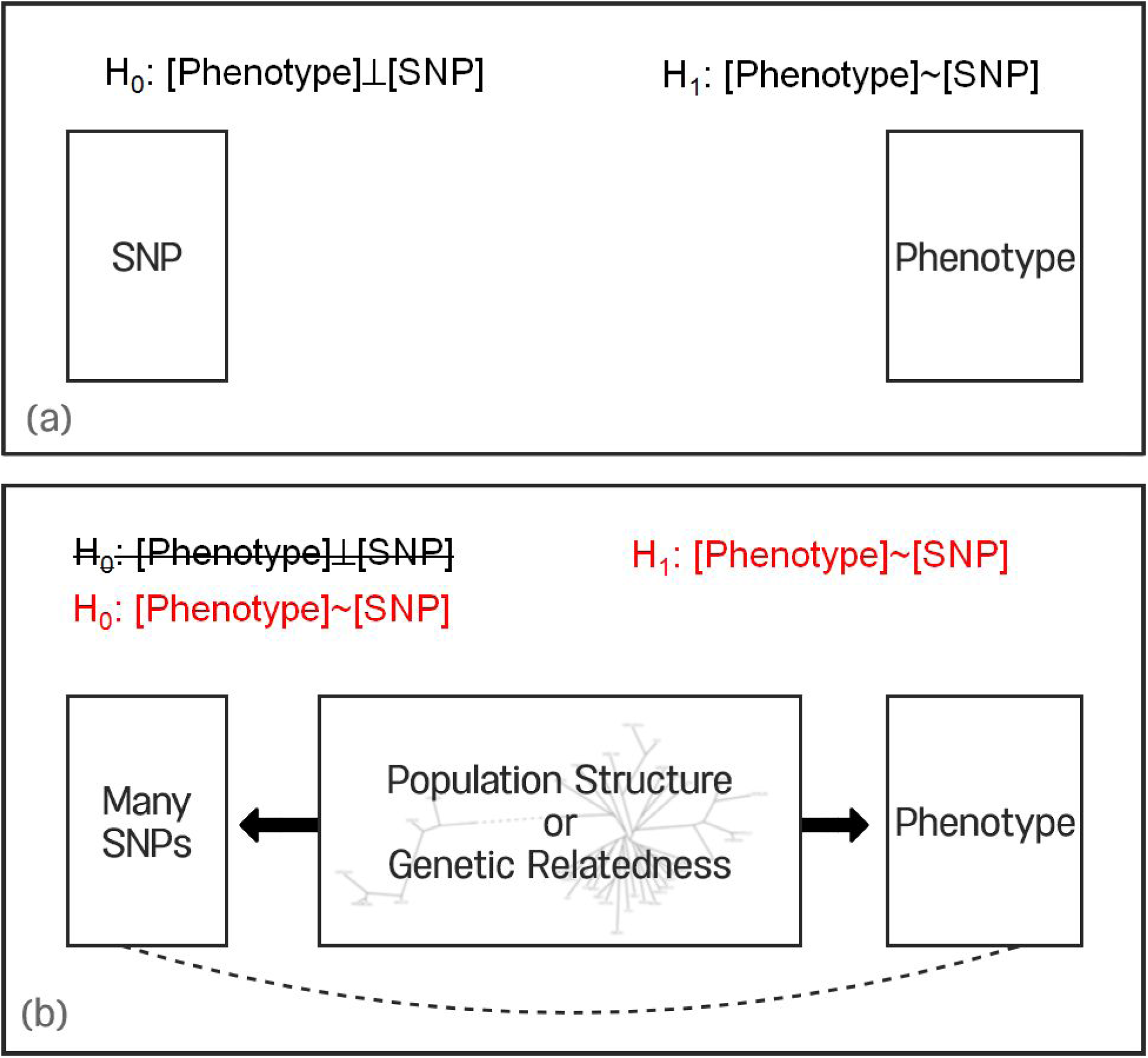
(a) The SNP and phenotype are independent under the null hypothesis (*H_0_*) and correlated under the alternative hypothesis (*H_1_*). (b) In the case of population structure, the structure will influence many SNPs and the phenotype. In this case, correlation between SNPs and the phenotype will be induced in both the null and alternate hypothesis.

The effects of these SNPs are the unmodeled factor in the equation shown in equation (3), and they confound our ability to perform association studies. In reality, there are many SNPs located on the long branching line (Figure 7, dashed line) that affect the phenotype. In order to identify these true associations, we must eliminate the unmodeled factor. While we cannot know which specific SNPs comprise the unmodeled factor, we can use available knowledge about similarities between the genomes of individuals in our studies to estimate the unmodeled factor.

**Figure 9.**
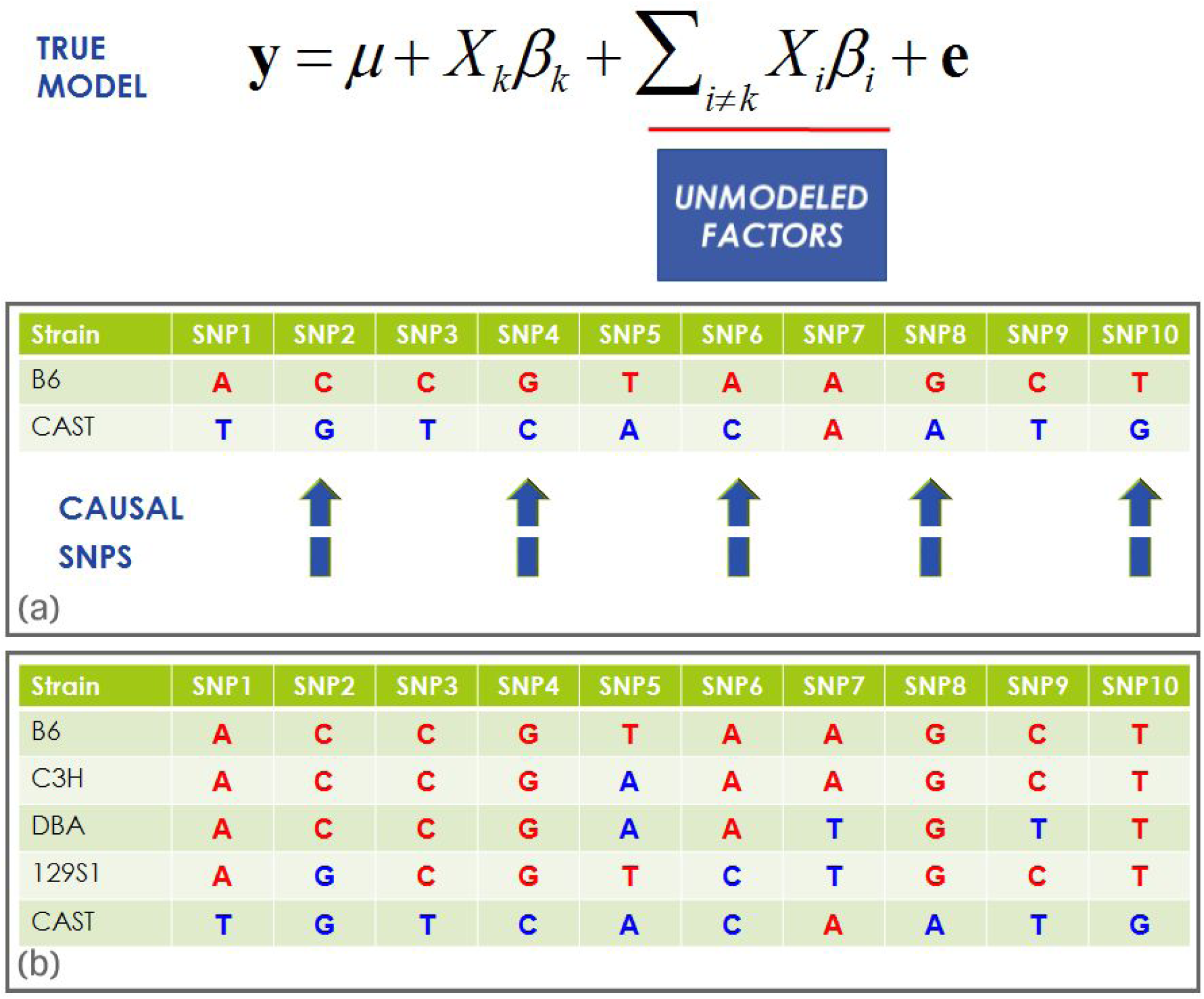
In a true model of association, the unmodeled factors cannot be known. We can estimate the unmodeled factors when (a) genomes are similar or (b) genomes share few causal variants.

Using our mouse example, we consider two different strains, B6 and C3H. These two strains are both classical inbred mice derived from domesticated mice and have similar genomes. In Figure 9a, we show a toy example considering the genomes of the two strains. Here, the genomes are very similar; nine out of ten SNPs are shared between B6 and C3H. In our example, let us assume that the even numbered SNPs are causal variants that affect the phenotype. For those variants, their corresponding effect size (*β_i_*) will be non-zero. We neither know the actual effect sizes nor the resulting value for the unmodeled factor. However, because they share the same allele as these SNPs, we do know that the two strains will have a similar value for the unmodeled factor.

Next, we consider two very different strains pairwise (Figure 9b): the classic inbred mouse strain B6 and the wild mouse strain CAST. In this case, the strains have different alleles present at many SNPs. If any of these SNPs affect the trait, the value of the unmodeled factor will differ by the effect size. Thus, we expect the two strains to have different values for the unmodeled factor.

The amount of pairwise sharing of alleles between strains can be used to capture the similarity between the values of the unmodeled factor among strains. In order to do this, we make a matrix that contains all SNPs shared between the paired genomes (Figure 10). This matrix allows us to “model” the values of the unmodeled factors among the individuals in our study, and it shows us which pairs have similar sharing of alleles and which pairs have dissimilar values.

The principle underlying mixed models is that we incorporate this “model” of unmodeled factors into the association test. We incorporate the unknown factors into the model of association using what is called a *random effect* or a *variance component*. Our model is called a *mixed model*, because it combines a random effect with the effect sizes of the SNPs we are testing (referred to as *fixed effects*) to model population structure.

**Figure 10.**
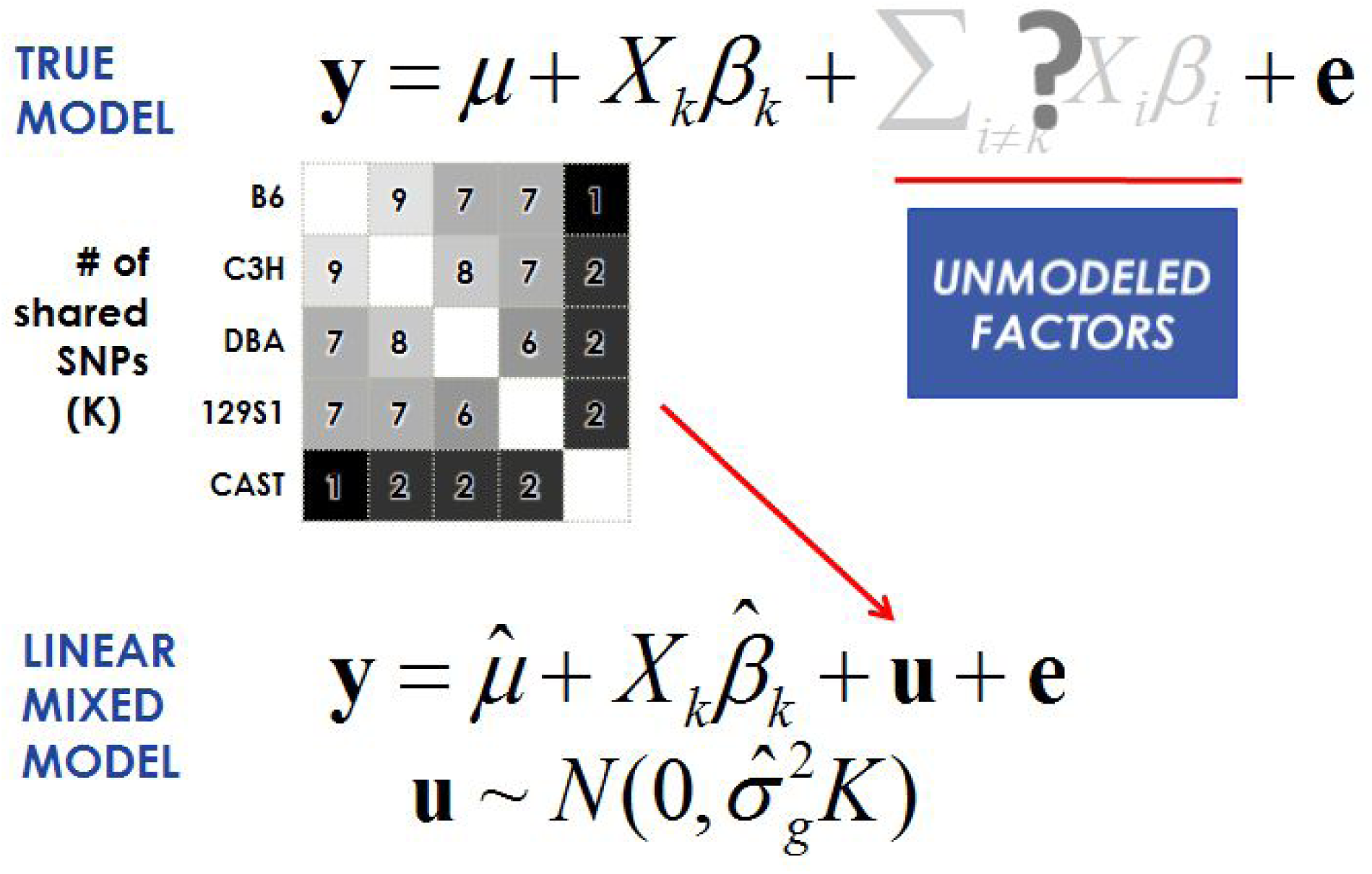
The true model ignores dependency among unmodeled factors and produces false associations. The mixed model reduces false associations by accounting for the dependency among SNPs correlated with phenotypes due to population structure.

When using a mixed model to identify causal variation, one key step is to establish these fixed parameters and random effect components. A linear mixed model (LMM) uses the information from the matrix to account for the unmodeled factor. We extend the simple, hypothetical true model

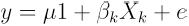

to include a term that captures the unmodeled factors. The term *u* in

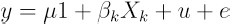

is a random vector that depends on the amount of shared genome in terms of pairwise differences. Here, we assume that *u*~*N*(0,*σ*^2^ *k*), where *K* is the kinship matrix. Each entry of *K* estimates the pairwise similarity between the genomes of the individuals in the study, which follows the intuition of Figures 9 and 10.

In practice, *K* can be computed from the genotypes where each entry in the kinship matrix is just the product of the standardized genotypes for the two individuals divided by the number of variants. Thus, the kinship entry computing the relatedness between individuals *i* and *j* is

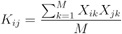

We can elegantly compute the kinship matrix using the equation *K* = *XX*^*T*^/*N*. The standard estimation equations above cannot be used to estimate the values of the parameters. Due to the random effect *u*, the phenotypes of the individuals are no longer independent of each other—an assumption of the previous methods.

However, if we know the values of 
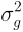
 and 
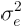
 we can then apply the following “mixed model trick.” We note that the phenotypes will follow the distribution

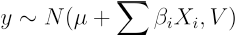

where 
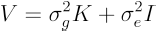
 and *I* is the identity matrix. If we transform then multiply the phenotypes and genotypes by 
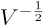
, we get

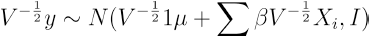

In the transformed data, the individuals are now independent of each other, and we can apply the estimation equations presented above to estimate the values for *β* and the association statistics.

In this case, we assume that the *β*_*i*_ values are drawn from a normal distribution with a mean zero as effect size and 
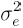
 as the variance.

Estimating the values of 
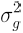
 and 
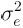
 is a difficult computational problem referred to as estimating the variance components. These parameters are estimated by utilizing a maximum likelihood approach. Specifically, we attempt to find the values of 
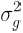
 and 
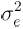
, such that the following log likelihood function of the data is maximized:

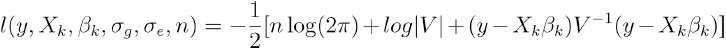

where 
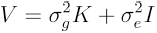
.

This equation is computationally difficult, because likelihood requires computing the inverse of the matrix (*V*^−1^), which in turn depends on the values of *σ*_*g*_ and *σ*_*e*_. Optimization methods that maximize this likelihood apply algorithms updating current estimates of of *σ*_*g*_ and *σ*_*e*_ until they converge to high values of the log likelihood function. Each step of an optimization algorithm is referred to as an iteration. In each iteration, the optimization algorithm must evaluate the log likelihood for the current values of *σ*_*g*_ and *σ*_*e*_ and must compute this matrix inverse. A straightforward way to compute a matrix inverse involves a complexity of approximately *O*(*n^3^*). Unfortunately, this results in a very inefficient algorithm and prevents mixed models from being widely utilized in association studies, despite their long history in genetics.

We developed Efficient Mixed Model Association (EMMA) (Kang et al. 2008), an efficient algorithm for estimating these parameters. Since we first presented EMMA, many other groups have developed similar efficient algorithms (Kang et al. 2010; Lippert et al. 2011; Zhou and Stephens 2012). The key idea behind EMMA is that we apply spectral decomposition to the kinship matrix, leading to a much faster optimization algorithm. The spectral decomposition only needs to be computed once and requires a complexity of *O*(*n*^3^). Specifically, if we write *K = UDU^T^* where *U* is a matrix of eigenvectors and *D* is a diagonal matrix of eigenvalues, then we can represent *V* using matrix algebra properties as follows:

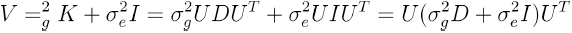

We can then compute the quantity 
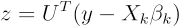
for each SNP *k* which has complexity *O*(*n^2^*). The log likelihood of the data can then be computed using

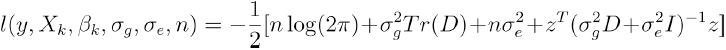

which can be computed in complexity *O*(*n*) since the matrix inside the likelihood is now diagonal. The inverse can be computed by simply taking the reciprocal of the elements along the diagonal. This procedure results in a very efficient algorithm that is useful for today’s large-scale human genomic datasets.

We applied EMMA to the same mouse association data analyzed using a standard LMM approach (see Figure 6). With these computational improvements, we almost completely reduced the inflation of false positives while obtaining nearly uniform p-value distribution for most SNPs (Figure 11). Here, the strongest peak, which is not significant, falls into a region of the genome on chromosome 8, which is known to be associated with body weight. Regions of the genome that correlate with variation in a phenotype are referred to as Quantitative Trait Loci (QTL).

Next, we applied EMMA to other phenotypes from the same mouse strain datasets, including a liver weight phenotype. Here, we see that the inflation of false positives is reduced and a strong signal at chr2 is more pronounced after the correction (Figure 12). EMMA correctly identifies a locus for liver weight that falls into the QTL Lvrq1 (liver weight), which was previously identified using a traditional mous mapping approach (Rocha et. al. 2004).

**Figure 11.**
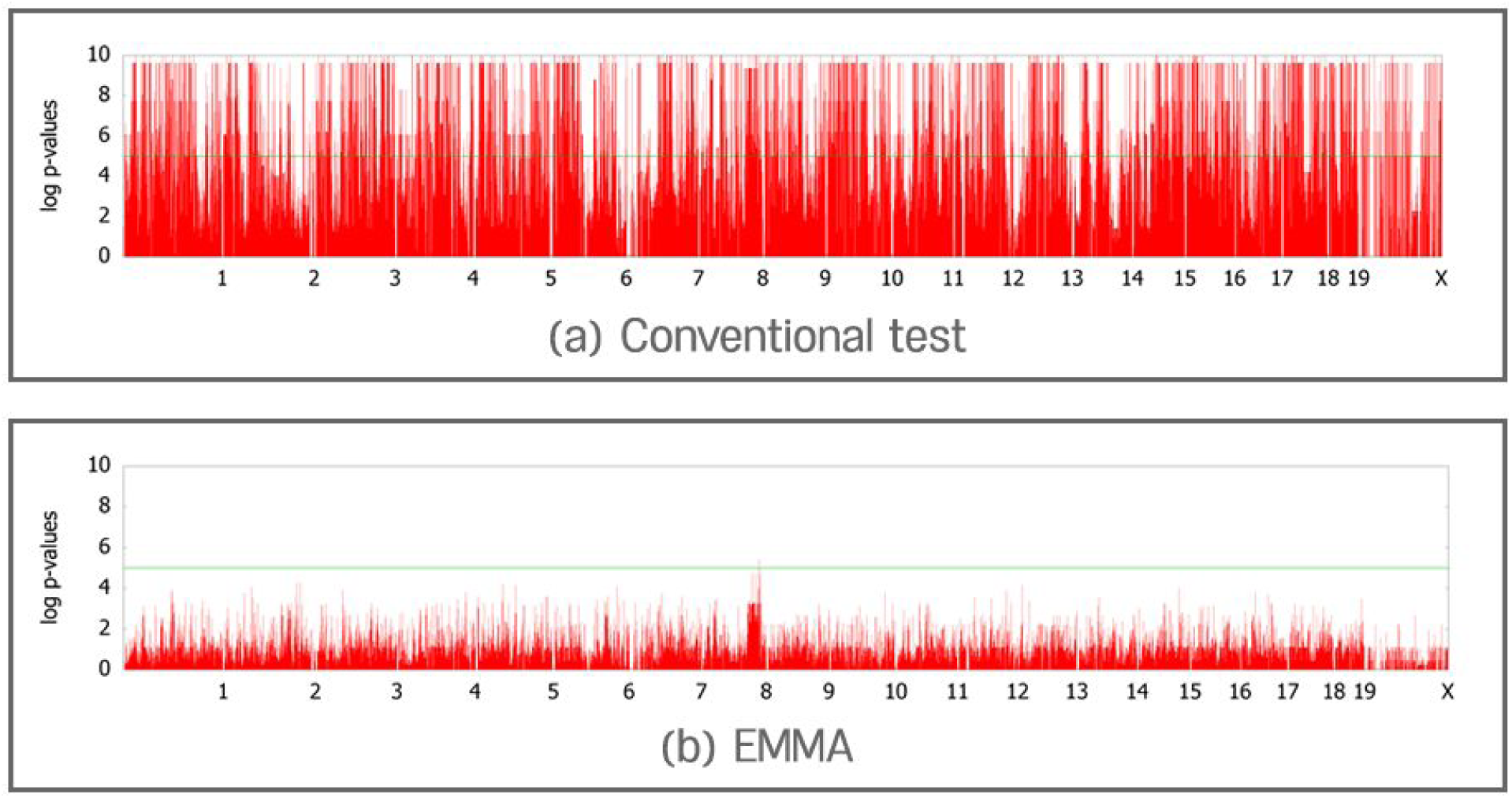
(a) The conventional GWAS test applied to mouse body weight phenotypes produces numerous false positions. (b) The mixed model approach using EMMA almost completely reduces the inflation of false positives and identifies a strong peak (chr8) that falls into a known body weight QTL.

**Figure 12.**
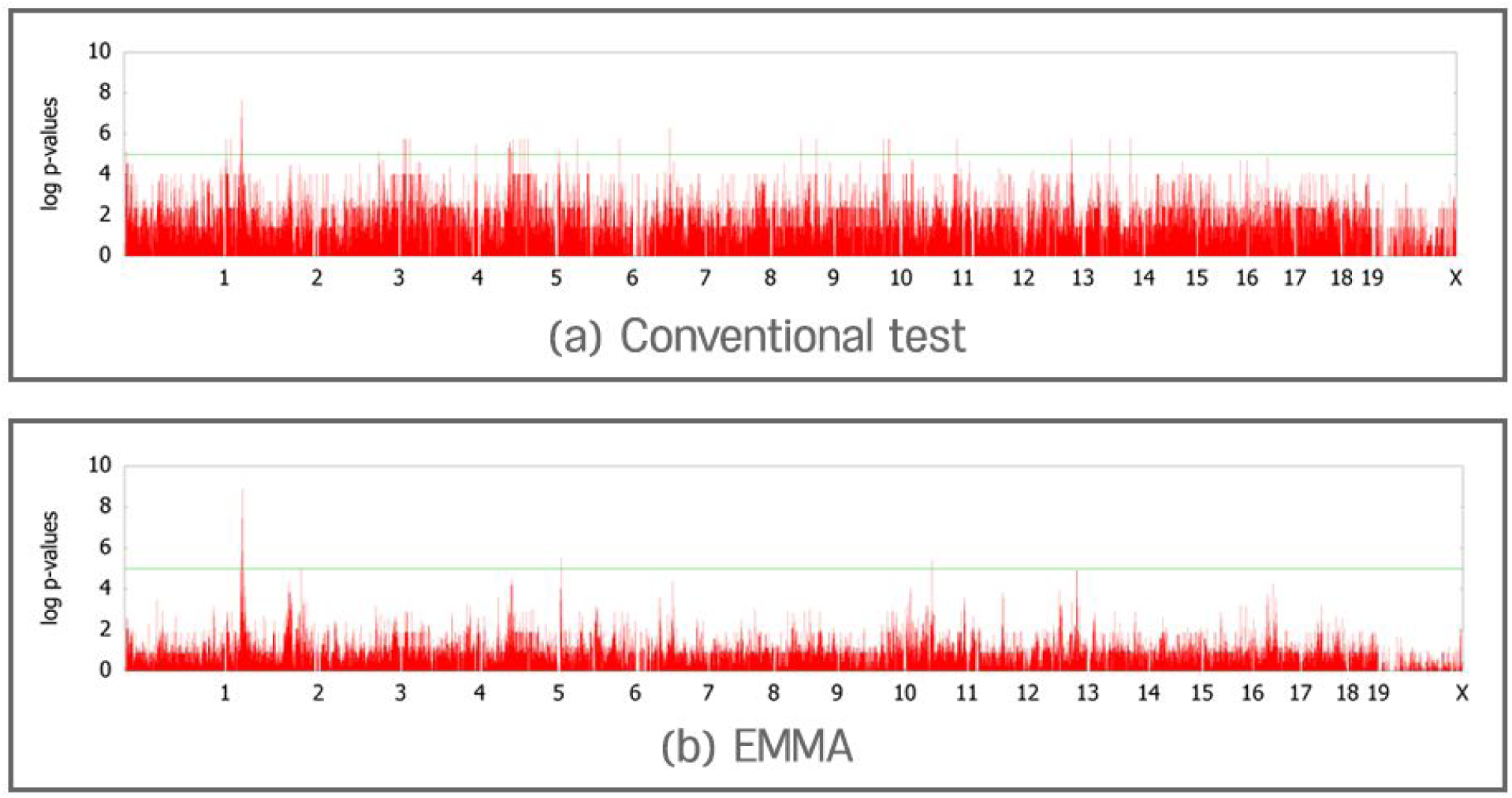
(a) The conventional GWAS test applied to mouse liver weight phenotypes produces numerous false positions. (b) The mixed model approach using EMMA reduces inflation of false positives and correctly produces a stronger signal at chr2, a region that is located in known QTLs for liver weight.

### Population Structure and Mixed Models in Human Association Studies

During the time that mixed models were starting to be used in mouse studies, the problem of relatedness in human studies was becoming apparent by causing difficulties in analyzing human GWAS studies. At that time, there was no single approach to handle relatedness. Instead, different types of relatedness were explicitly modeled, and association study methods were adapted to those scenarios. There is an entire class of methods designed to handle relatedness when there are closely related individuals in the genetic study and the genetic relationships are known. These include methods for multigenerational families, twins, and siblings (Freimer and Sabatti 2004; Van Dongen et al. 2012)

A complication in human association studies is when the relationships are unknown. One of the most common types of relatedness among individuals in human studies is due to ancestry. *Ancestry* refers to the population that an individual descended from. Many individuals are admixed, which means they are descended from ancestors in different populations. If an association study contains individuals from different populations or differing degrees of admixture, the individual will have different degrees of relatedness among them. In other words, individuals with the same ancestry are slightly more related to each other than individuals with different ancestries.

It is well documented that these ancestry differences can induce false positive associations (Helgason et al. 2005). Association studies that analyzed individuals with differences in ancestry typically utilized an approach to predict the ancestry for each individual and then incorporated this information as a covariate in the model (Pritchard et al. 2000). An alternate approach was to estimate principal components over the genotype data, which could be interpreted as a proxy for association studies and included in the model as covariates (Price et al. 2006). In the human genetics literature, ancestry differences are sometimes referred to as population structure. In this review, we use the term *ancestry differences* separately from the term *population structure*; we use the latter to describe the general phenomenon of relatedness in a sample.

A second type of relatedness is cryptic relatedness (Voight and Pritchard 2005). Since GWAS are applied to extremely large samples, there are often individuals included in the study who happen to be related—but this relatedness is unknown the both the individuals and the investigators. Typically, cryptic relatedness is handled by screening the association study for related individuals and computing the genetic similarity between each pair of individuals.

A general purpose approach to correct for population structure, or any type of confounding in association studies, is genomic control (Devlin and Roeder 1999; Bacanu et al. 2002). Genomic control allows us to measure the extent to which population structure (or other confounders) is affecting the association statistics. By examining the cumulative p-value distribution plot, we consider the deviation of the actual plot from what is expected at the median. Since we expect the vast majority of variants not to be associated with the trait, we expect the median observed p-value to be close to 0.5. Typically, population structure induces a more significant observed median p-value.

Genomic control computes a correction factor referred to as *λ*, which is a scaling factor used to scale all of the observed p-values so that the corrected median p-value is then 0.5. The *λ* is on the *χ^2^* scale (meaning that the median p-value is converted to a *χ^2^* value and the ratio is computed relative to the *χ^2^* value) corresponding to a p-value of 0.5, which is 0.545. The observed association p-values are converted from p-values to *χ^2^* statistics, scaled by *λ* and then converted back to p-values.

We can also use the value of the *λ* as a measure of the extent of the effect of confounding on the association statistics. Genomic control *λ*’s are widely utilized to compare different correction approaches. A *λ* of 1.0 shows that there is no inflation. A value greater than 1.0 is evidence that the association statistics are inflated. Typically, the 95% confidence interval of the *λ* in GWAS studies is 0.02. Thus, any *λ* of 1.03 or higher suggests that there is some inflation. We note that more recent exploration of polygenicity, or the amount of causal variants for a trait, suggests that there are many more causal variants than originally expected. In this case, the *λ* values should actually be higher than 1.0 (Yang et al. 2011). We discuss this perspective in the Discussion.

In the literature, ancestry differences and cryptic relatedness are referred to as distinct phenomenon. In fact, they can be thought of as different degrees of relatedness in the sample. Consider in Figure 13a, which shows a potential pedigree relating all of the individuals in an association study sample. Ancestry differences can be thought of relatedness near the top of the tree (Figure 13b), and cryptic relatedness can be thought of relatedness in a more recent portion of the tree (Figure 13c).

**Figure 13.**
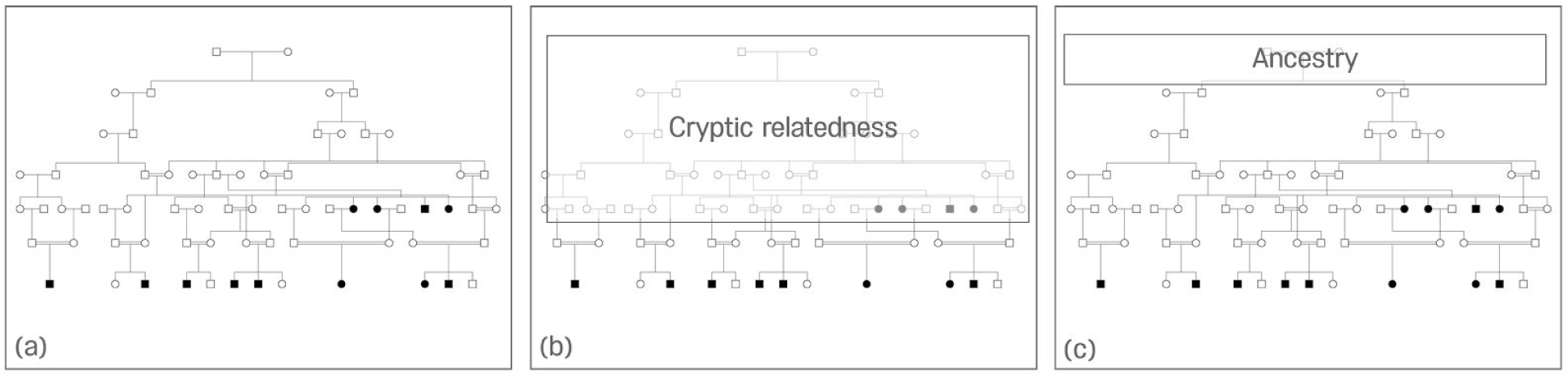
Different degrees of relatedness in the sample. (a) All of the individuals in a genetic study are somehow related through a large pedigree or family tree. This tree can produce two forms of hidden relatedness: cryptic relatedness (b) and ancestry (c), where the box represents the level of the pedigree that causes that type of relatedness.

Mixed models can handle nearly arbitrary genetic relationships between individuals and are a natural approach for human association studies. Mixed models are ideal because they can be applied without explicit identification of the ancestry and relatedness within the sample. They also enable the analysis of datasets with particularly complex genetic relationships, such as isolate populations where the population is descended from a small number of founder individuals (Kenny et al. 2010). For isolate populations, the previous methods were not able to fully account for population structure.

Mixed models were first used in human studies with the Northern Finnish Birth Cohort (Sabatti et al. 2009), where mixed models were applied to 331,475 SNPs in 5,326 individuals who were phenotypes for 10 traits (Kang et al. 2010). These traits include C-reactive protein (CRP), triglyceride (TG), insulin plasma levels, (INS), diastolic blood pressure (DBP), body mass index (BMI), glucose (GLU), high-density lipoprotein (HDL), systolic blood pressure (SBP), and low density lipoprotein A (LDL). Individuals within this cohort have some ancestry differences due to their origin from different parts of Finland, and they share some genetic relationships.

Table 1 shows the results of applying mixed models to these traits. Each entry in the table shows the *λ* value for the analysis of that phenotype. The first column shows the results of the uncorrected analysis. We can see that there are very large *λ* factors, particularly for height. In fact, the associations with height were not reported in the original Sabatti et al. (2009) manuscript because the high *λ* value suggested that some of the observed associations may be false positives. The second column shows the *λ* factors after eliminating cryptically related individuals. Here, we compute the pairwise relationships between individuals and filter out one of any pair that was closely related. This approach filtered out 611 individuals.

The third column shows the *λ* factors after using 100 principal components as covariates. This was done to show the limit of the principal component approach in correcting for population structure. Each component decreases the *λ*; using 100 components is an absurdly large number of components and is well beyond what is typically utilized in any type of association study. The last column shows the *λ* for mixed models. Each of these *λ* values are within the 95% confidence interval (around 1.0), suggesting that mixed models can correct for all of the population structure in the sample—including cryptic relatedness and ancestry differences. As shown in Table 1, only mixed models adequately correct for population structure in this sample.

**Table 1.** Results of analysis (*λ* values) on NFBC66 data.

| Traits | Uncorrected | IBD<0.1 | 100PC | EMMAX |
| --- | --- | --- | --- | --- |
| BMI | 1.036 | 1.028 | 1.024 | 1.001 |
| CRP | 1.012 | 1.020 | 1.020 | 0.994 |
| DBP | 1.033 | 1.025 | 1.029 | 1.010 |
| GLU | 1.045 | 1.025 | 1.030 | 1.009 |
| HDL | 1.054 | 1.041 | 1.037 | 1.003 |
| INS | 1.026 | 1.026 | 1.015 | 1.005 |
| LDL | 1.093 | 1.089 | 1.040 | 1.002 |
| SBP | 1.063 | 1.054 | 1.021 | 1.004 |
| TG | 1.024 | 1.021 | 1.018 | 0.999 |
| HEIGHT | 1.193 | 1.152 | 1.080 | 1.002 |

Mixed models have become important in human GWAS analysis, because the estimates of 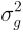
 and 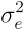
 can be used to estimate the heritability of the trait. Recent results suggest that common variants explain a larger proportion of the variance of complex traits than previously thought (Purcell et al. 2009; Yang et al. 2010; Eskin 2015).

## Discussion

Over the past decade, association studies have identified thousands of variants implicated in dozens of common human diseases. The traditional approach to association studies assumes that individuals are unrelated to each other. However, in practice, individuals in genetic studies are related to each other in complex ways. In this review, we demonstrate how these relationships cause false positives in association studies and how mixed models can correct for these confounding genetic relationships.

This review covers only the basic principles of mixed models and population structure. Since the original EMMA paper in 2008, mixed models have become an active research area. Many groups have published papers exploring various aspects of mixed models and their application to complex genomic problems.

For example, many approaches have been developed to improve the efficiency of mixed models, including the methods Fast-LMM (Lippert et al. 2011) and GEMMA (Zhou and Stephens 2012). More recently, a method called BOLT-LMM (Loh et al. 2015) was developed for scaling analyses to handle cohorts in the hundreds of thousands of individuals.

Another direction of method development has been extending mixed models to handle case control studies. These approaches typically assume a liability threshold model where there is an underlying continuous phenotype; if the phenotype is above a threshold, the individual has a disease. If it is below a threshold, the individual does not have the disease (Zaitlen et al. 2012). These types of studies are also complicated by a phenomenon of selection bias, because the cases are oversampled from the population. At present, such mixed model extensions to case/control studies results in challenging computational problems (Hayeck et al. 2015; Weissbrod et al., 2015).

Some mixed models are developed based on observation of a particular bias inherent to standard approaches. For example, a bias is induced by the SNP that is tested and used in the computation of the kinship matrices (Listgarten et al. 2012). This bias motivated the idea that, when applying mixed models, the kinship matrix should not contain the SNP being tested. As a result, the Leave One Chromosome Out (LOCO) approach constructs a different kinship matrix for testing each chromosome and leaves out the SNPs on the chromosome being tested (Yang et al. 2014).

This approach is also motivated by the observation that many complex traits are highly polygenic, suggesting that there are hundreds (if not thousands) of loci that influence some traits (Yang et al. 2011). Some traits, such as height, are known to be highly polygenic. In this case, it is not clear what the actual value of *λ* should be for a polygenic trait as it is expected to have a contribution from both confounding effects as well as polygenicity. More recently, a method called LD score regression has been developed that attempts to differentiate between these two components (Bulik-Sullivan et al. 2015).

From their origins in non-human organisms to powering large scale human genome wide association studies today, mixed models play an important role in the analysis of genetic data, particularly in correcting for population structure. Research in improving and extending mixed model approaches is now an active research area in the field.

